# Testing for context-dependent effects of maternal thyroid hormones on offspring survival and physiology: an experimental approach manipulating temperature in a wild bird species

**DOI:** 10.1101/2020.05.08.083725

**Authors:** Bin-Yan Hsu, Tom Sarraude, Nina Cossin-Sevrin, Melanie Crombecque, Antoine Stier, Suvi Ruuskanen

## Abstract

Maternal effects via hormonal transfer from the mother to the offspring provide a tool to translate environmental cues to the offspring. Experimental manipulations of maternally transferred hormones have yielded increasingly contradictory results, which may be explained by environment-dependent effects of hormones. Yet context-dependent effects have rarely been experimentally tested. We therefore studied whether maternally transferred thyroid hormones (THs) exert context-dependent effects on offspring survival and physiology by manipulating both egg TH levels and post-hatching nest temperature in wild pied flycatchers (*Ficedula hypoleuca*) using a full factorial design. We found no clear evidence for context-dependent effects of prenatal THs related to postnatal temperature on growth, survival and potential underlying physiological responses (plasma TH levels, oxidative stress and mitochondrial density). We conclude that future studies should test for other key environmental conditions, such as food availability, to understand potential context-dependent effects of maternally transmitted hormones on offspring, and their role in adapting to changing environments.

## Introduction

Maternal effects may translate environmental cues from the mother to the offspring for example via maternally transferred hormones (hereafter ‘maternal hormones’), potentially increasing offspring survival in the predicted conditions (adaptive maternal effects, 1, 2–4). Maternal hormone-mediated effects have been recently highlighted as a potential mechanism and source of phenotypic plasticity to respond to changing climate (5–7), yet empirical evidence is scarce. Experimental studies on hormone-mediated maternal effects have revealed increasingly contradictory results, for example, elevated maternal androgens both increasing, decreasing or having no effect on offspring growth (3). One obvious explanation for such contrasting effects is linked to the well-known pleiotropic effects of (maternal) hormones, due to which prenatal hormone exposure brings both costs (*e.g*. reduced immune response) and benefits (*e.g*. faster growth). Therefore, the final fitness outcome is likely to be determined by the potential benefit-cost balance being set by environmental conditions (3).

Studies on context-dependent effects of maternal hormones have been repeatedly called for in the literature, but still only few empirical studies are available. In the most studied maternal hormone group, glucocorticoids, fitness effects depend on the matching between maternal and offspring environment (e.g. 8, 9), such as maternal condition (*e.g*. (10, 11), density (12) and predictability of food (13). In the rare example on maternal androgen hormones, Muriel et al. (14) suggest that the effects of maternal androgens on nestling development and immunity depend on the timing (first or second clutch) of breeding and associated food conditions. To our knowledge, the context-dependent effects of other key maternal hormones, such as thyroid hormones, have not been explored. Yet, characterizing context-dependent effects can contribute to our understanding on the cause of the observed large variation in maternal hormone transfer across and within populations (4, 15) and its potential adaptive function.

We previously reported contrasting effects of maternal thyroid hormones (THs: T3=triiodothyronine, T4=thyroxine) on offspring phenotype in two closely related species, the collared flycatcher (*Ficedula albicollis*) and pied flycatcher (*F. hypoleuca*) (16, 17). THs increased early growth, but decreased growth during the second week post-hatching in collared flycatchers (16), while they tended to increase growth during the second week post-hatching in pied flycatchers (17). The underlying mechanisms and the explanations for these contrasting results remain unknown, but one hypothesis could be that the effects of elevated yolk THs would depend on the post-hatching environmental conditions. For example, if prenatal THs increase resting metabolic rate (RMR), as reported in Hsu et al. (18), the elevated RMR may lead to increased growth in benign conditions, but decreased growth when resource availability is poor (19). Moreover, THs have conserved function in controlling thermoregulation (20), which is not fully developed in altricial offspring until late postnatal stage (21). If prenatal THs are expected to increase metabolic rates and stimulate thermogenesis, they may benefit nestling survival in low developmental temperatures, while in higher temperatures such effects may not be observable or even turn negative because of higher energy expenditure. Therefore, experimental tests on whether the effects of yolk THs would depend on post-hatching environmental conditions are now duly required.

Here, we use a full factorial experimental design to, for the first time, study whether maternal THs exert context-dependent effects on early-life phenotype, survival and physiology. We experimentally manipulated both egg TH levels and post-hatching nest-box temperature in a wild population of pied flycatchers. We chose to study interactions between THs and temperature variation, because both factors are involved in thermoregulation (7, 20), which is crucial for early-life altricial nestlings (21). Moreover, we previously found that yolk T4 are higher under relatively lower ambient temperature in passerines (22). Testing whether yolk THs have temperature-dependent effects therefore would provide information on whether elevated T4 transfer under lower temperatures could be an adaptive allocation. In addition to measuring postnatal growth and short-term survival (fledging success), we explored changes in potential underlying physiological mechanisms. (i) We measured circulating THs to track any lasting effects of prenatal TH manipulation on the general function of the hormonal axis (18) and to assess the direct effects of postnatal temperature on THs. (ii) We estimated mitochondrial density (*i.e*. mitochondrial DNA copy number) as a proxy for the effects of our treatments on cellular bioenergetics (23). Mitochondria are the powerhouse of cells, converting nutrients into ATP to sustain cellular functions. Mitochondrial density and bioenergetics are likely to be influenced by THs across taxa (24, 25, reviewed in 26) and by ambient temperature (27). However, to our knowledge, the effects of prenatal THs on mitochondria have not been characterized beyond mammalian models (28). Finally, we measured (iii) biomarkers of oxidative stress, the imbalance between oxidizing molecules (*e.g*. reactive oxygen species, ROS) and antioxidant protection, which ultimately leads to oxidative damage on biomolecules and cellular dysfunction (29). Elevated yolk THs may result in increased oxidative stress directly via the stimulating effects of THs on metabolism and mitochondrial ROS production (30), or indirectly via increasing developmental speed since fast growth is likely to increase oxidative stress (31). Temperature is also likely to influence oxidative stress since both cold and heat stress have been shown to increase oxidative damage levels (32, e.g. 33).

We summarize our predictions in Table 1. Overall, we predicted that elevated prenatal THs should increase survival in cooler (non-heated) nests, for example due to its stimulating and positive effects on thermogenesis, which might also facilitate growth. On the other side, the increased metabolism in nestlings from heated nests might lead to higher energy expenditure, which albeit not necessarily decreasing survival, but might reduce growth (e.g. 18). We predict that elevated prenatal THs increase postnatal circulating THs and the effect may be stronger in cooler (non-heated) nests. Mitochondrial density is predicted to be increased by the stimulating effect of (prenatal) THs and low temperature, and thus being highest in the nestlings from TH-injected eggs and non-heated nests. Finally, oxidative damage is also predicted to be higher in prenatal TH elevation group (see above), which could be more pronounced in heated nests due to maladaptive thermogenesis enhancement.

**Table 1.** Summary of the predictions regarding changes in the response variables in relation to elevated prenatal thyroid hormones (THs), postnatal heating and their interaction, i.e. context-dependent effects. + = positive effect of TH elevation/heating, - = negative effect of TH elevation/heating.

| **Nestling trait** | **Prenatal TH elevation** | **Postnatal heating** | **TH x Heating interaction** |
| --- | --- | --- | --- |
| **Survival** | NA | NA | Elevated prenatal TH increases nestling survival in non-heated nests (due to stimulating effects on thermoregulation) |
| **Growth** | ? | ? | Elevated prenatal THs decrease nestling growth in heated nests (unnecessarily elevated metabolism caused higher energy expenditure) |
| **THs** | + | - | Highest in TH+non-heated, lowest in CO-heated |
| **Mt density** | + | - | Highest in TH+non-heated, lowest in CO-heated |
| **Oxidative damage** | + | 0/- (low magnitude) | Highest in TH+heated, lowest in CO+heated |

## Material and methods

The experiments were conducted under licenses from the Animal Experiment Board of the Administrative Agency of South Finland (ESAVI/2902/2018) and the Environmental Center of Southwestern Finland (license number VARELY549/2018). We conducted our experiment in a nest-box population of pied flycatchers located in Turku, Finland (60°25’N, 22°10’E). Egg THs were manipulated in unincubated eggs using injections in the yolk, following (17, 34). All eggs of a nest received the same injection of either thyroid hormones (hereafter, TH clutches, N = 30 clutches) or only vehicle (hereafter, CO clutches, N = 30 clutches). Hatching success was not affected by the hormone treatment (CO: 71.1%, TH: 64.9 %, z = −1.129, p = 0.259).

On the second day after hatching (d2), chicks were individually identified by nail clipping and *ca*. half of the chicks of each nest were swapped among broods with different hormone treatments whenever possible, to create nests which included both CO and TH treated nestlings. Internal temperature of the nest boxes was then increased between postnatal day 2 and day 8 in *ca*. half of the boxes (hereafter, ‘heated nests’, N = 24) using heating pads (UniHeat 72h, USA), installed under the ceiling, and replaced every 2^nd^ day. The period of 2 to 8 days was selected as pied flycatcher nestlings are not fully thermoregulatory during this period, and thus sensitive to variation in ambient temperature (35). The other half served as controls (hereafter ‘non-heated nests’, N = 23 nests) and received non-functional heating pads and similar visits every 2^nd^ day. The sample sizes in the four treatment groups at day 2 are: CO+non-heated 76, CO+heated 50, TH+non-heated 58, and TH+heated 53 nestlings, respectively. The actual temperature within the boxes was recorded with a thermo-logger (iButton thermochron, measuring at 3min intervals, 0.0625°C accuracy), placed inside all nest boxes at 5 cm distance above the nest rim, and the average daily temperature between d2 and d8 was calculated. The heating treatment increased the temperature by 2.75 ºC (SE=0.37, marginal means±SEs controlled for date and iButton position: heated 21.25±0.36, non-heated 18.50±0.35, GLM, t=-7.40, p<0.001, Fig. S1).

Female can be actively brooding until the nestlings are *ca*. 6-7 days old (35). To account for potential behavioral changes of the female linked to the heating treatment that could influence offspring traits (potentially mask any effect of the heating treatment), we collected data on brooding behavior on d4 after hatching. We recorded the percentage of time spent brooding during a minimum of 2 hours, using miniature video cameras (ca 4×4cm, DashCam, UK) mounted at *ca*. 2 m distance from the nest-boxes (N_non-heated_ = 16, N_heated_ = 13). Females brooded *ca*. 33% of the time, but the variation was large (SD 16%) and there was no significant difference in brooding behavior between the treatments (mean ± SD: non-heated nests 34.7±16.3%; heated nests 36.3±15.8%; t = 0.58, p = 0.57), suggesting that any potential effect of the heating treatment would not be influenced by parental behavior. This data also suggests that during daytime, offspring are exposed to ambient temperatures for substantial periods of time (60% of the time at d4).

Nestling survival was checked at every nest visit (d2, d4, d6, d8, d13). Nestling body mass (~0.01g) was recorded at d2, d8 and d13 after hatching, and tarsus length (~0.01 mm, proxy for skeletal growth) at d8 and d13 after hatching. A blood sample (*ca*. 40μl) from the brachial vein was collected using heparinized capillaries at d13 after hatching and centrifuged before being frozen at −80°C. Plasma was used for measuring thyroid hormone concentration (see below). Blood cell pellets were used for extracting DNA to molecularly sex the nestlings and assess mitochondrial density (see ESM for details). Another sample of *ca*. 20-30 μl whole blood was collected directly in liquid nitrogen and thereafter stored at −80°C to analyze oxidative stress biomarkers.

Plasma T3 and T4 levels were analyzed using nano-LC-MS/MS following previously published methods (36) and are expressed as ng/ml. Two randomly picked samples per nest were selected due to logistical constraints. Two oxidative stress biomarkers, (i) oxidative damage to lipids (malonaldehyde, MDA, nmol/mg protein, using TBARS assay) and (ii) total glutathione (hereafter tGSH; μmol GSH/mg protein), the most abundant endogenous intracellular antioxidant, were measured using established protocols, following Sarraude et al. (17). For oxidative stress biomarkers, we excluded the nests whose nestlings were not involved in cross-fostering and the numbers of nestlings analyzed were 179 and 155 at d8 and d13, respectively. Mitochondrial density, estimated through relative mitochondrial DNA copy number, and molecular sexing were analyzed using qPCR on all nestlings survived to d13 (n=185), following methods in Stier et al. (23) and (37). See ESM for details.

### Statistical analysis

All statistical models were conducted in the environment of R 3.5.1 (R core team 2018). We ran separate generalized linear mixed models (GLMMs, package lme4 and pbkrtest, 38, 39) for each trait of interest to assess the interaction between the treatments of yolk TH injection and nest-box heating, in the meanwhile controlling for relevant covariates and random intercepts to account for potential non-independence among nestlings. The final sample sizes varied by traits due to both the logistical constraints (see above) and missing values due to random failures in certain assays. The exact sample sizes and model details are reported in ESM. For fledging success, binomial distribution and logit link function were applied. For blood MDA, tGSH, and mtDNA density, data were first ln-transformed, and for all models examined, no clear violation on residual normality was visually detected. We found no indications of sex-dependent effects of THs or temperature treatment for any of the response variables (all F < 1.5, p > 0.2), and thus those are not discussed further.

## Results and Discussion

To our knowledge, we present the first experimental study on context-dependent effects of maternally transferred THs. We found no clear evidence for temperature-dependent effects of maternal THs on offspring survival (fledging success, CO-non-heated 69.7% CO-heated 78.0% TH-non-heated 75.9% TH-heated 76.9%, z = 0.156, p =0.876, Table S1). These results suggest that contrary to our predictions, elevated prenatal THs do not clearly benefit nestlings’ survival in colder environment. Egg TH concentrations were previously observed to increase with decreasing ambient temperature in passerines (22). If that was an adaptive allocation, we would expect elevated yolk THs to benefit offspring survival under colder condition, but our results do not provide clear support for such an adaptive explanation.

The growth of offspring (tarsus or body mass) from TH elevated and control eggs was not differentially affected by the postnatal temperature treatments at any of the measured age-points (All F <1.30, p >0.26, Fig 1a, Table S2, S3). Our results therefore suggest that the previously reported discrepancies in the effects of prenatal THs on offspring growth across sister species (collared and pied flycatchers, 16, 17), and other altricial birds (18, 40) are unlikely driven by context-dependent effects of THs related to early postnatal temperature differences. Yet, there are several possible alternative explanations: (i) The study took place in a relatively warm year, with the average temperature during the nestling phase (June-July) being *ca*. 2 ºC higher than the averages in the past 15 years (2018: +18.4 º C vs 2003-2018 +16.5 º C). We may expect that the differential effects of THs on growth to be evident only under relatively cold conditions. (ii) Our experimental temperature manipulation was rather small, which could have either been compensated by maternal brooding (though brooding did not seem to differ among the groups) or simply too small to induce measurable changes in growth. In previous studies, ca 5 ºC elevation led to a decrease in body mass in blue tits (*Cyanistes caeruleus*, 33) and great tits (*Parus major*, 41, 42) and an increase in body mass in tree swallows (*Tachycineta bicolor*, 43); (iii) If maternal hormone transfer varies according to environmental context, our egg injection treatment might have resulted in different doses depending on the initial yolk hormone levels. Quantifying egg hormonal content in unmanipulated eggs across experiments, years and contexts is now needed to test this possibility. The logical next steps to test other context-dependent effects of maternal THs would be either to reduce ambient temperature during postnatal development or to manipulate some other environmental factors, such as food availability following similar experimental set-ups.

**Fig. 1:**
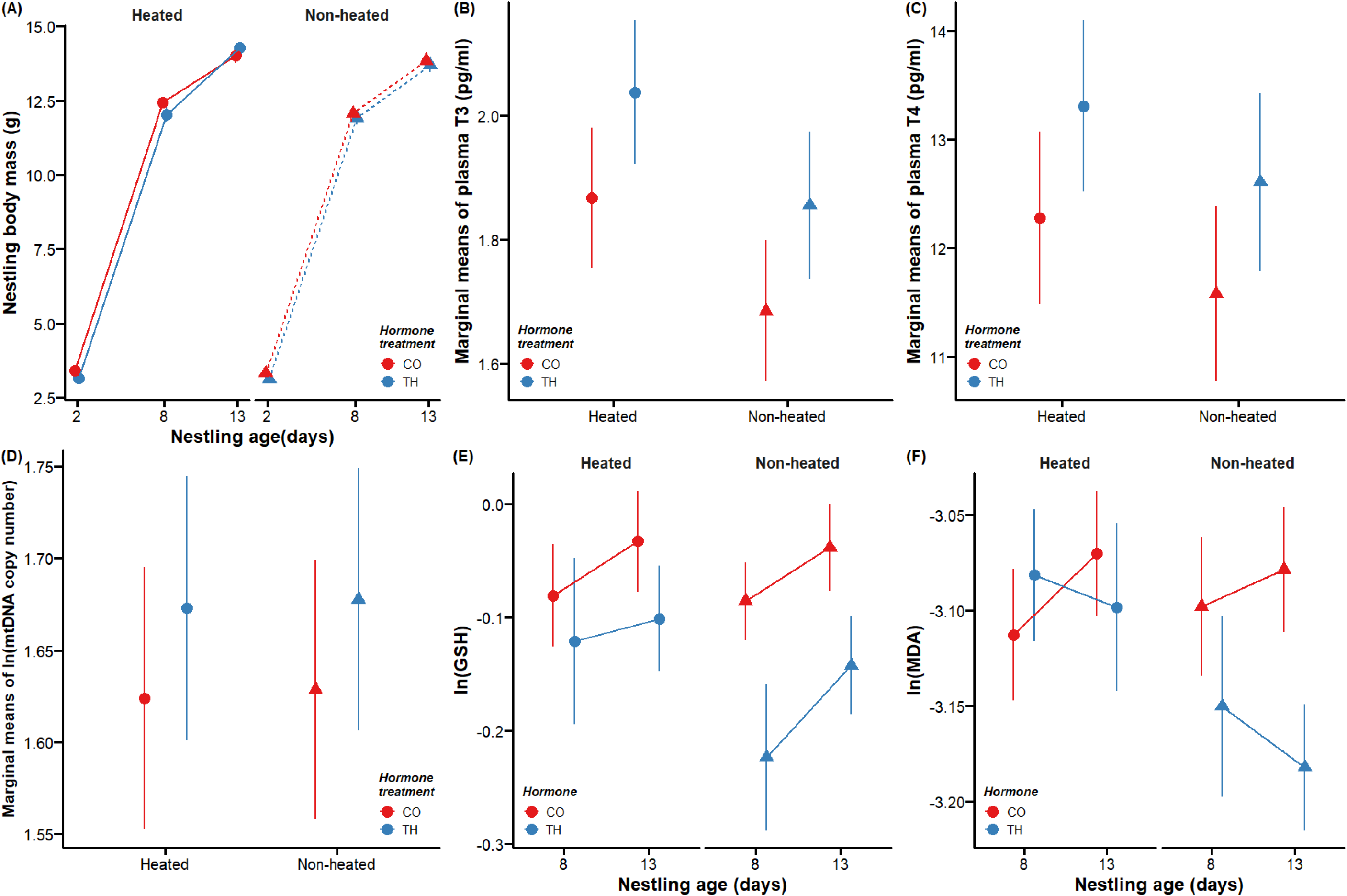
Effects of prenatal hormone manipulation (TH= experimentally elevated yolk thyroid hormone treatment, CO = control) and postnatal temperature manipulation (non-heated *vs*. heated nests) on offspring phenotype and physiology. (A) Nestling body mass growth pattern (g, average ± SE); (B) plasma triiodothyronine (T3) concentration (pg/ml, marginal means ± SE), (C) plasma thyroxine (T4) concentration (pg/ml, marginal means ± SE), (D) mitochondrial density in blood cells (ln-transformed, marginal means ± SE), (E) blood total glutathione concentration (tGSH, nmol/mg protein, ln-transformed means ± SE) and (F) lipid peroxidation (MDA concentration, μmol/mg protein, ln-transformed means ± SE). Heated nests were on average 2.75ºC warmer than non-heated ones. See text and ESM for details on statistics and sample sizes.

We aimed to characterize the nestling physiological changes underlying potential context-dependent effects of prenatal THs in response to variation in postnatal ambient temperature. Physiological biomarkers are commonly used as early proxies of responses to environmental variation, and are often sensitive even in cases where effects on growth and (early-life) survival are not visible (e.g. 44, 45). In contrast to our predictions, we observed no apparent differences in the circulating T3 or T4 concentration in nestlings exposed to higher levels of THs prenatally (T3: F_1, 28.3_ = 1.553, p = 0.223; T4: F_1,34.5_ = 1.032, p = 0.317), or in interaction with postnatal temperature treatments (hormone × heating, T3: F_1. 37.3_ =1.896, p = 0.177; T4: F_1,32.1_ = 0.028, p = 0.867, Fig. 1b,c, Table S4). Yet both T3 and T4 levels correlated positively with nestling body mass (estimate ±SE, T3: 0.196±0.035, F_1.36.5_ = 26.9, p < 0.001; T4: 0.563±0.219, F_1,33.2_ = 5.65, p=0.023, Table S4). Maternal hormones are suggested to cause long-lasting effects on offspring via changes in the function and sensitivity of the corresponding hormonal axis (3), so-called organizational effects. Changes in hypothalamus-pituitary-thyroid (HPT)-axis in response to prenatal THs during embryonic development has been characterized in chicken (46), pigeons (18) as well as mammalian models (reviewed in 47). Our results suggest an absence of such long-term programming effects of the HPT-axis in our study system – however it must be noted that circulating TH levels are highly variable in response to internal and external (food, temperature, circadian) variations (4, 46), which could mask potential organizing effects. Experimental challenges with thyrotropin-releasing hormones (TRH) or thyriod-stimulating hormone (TSH) are now needed to test for the effects of prenatal THs on the sensitivity of the HPT-axis in birds.

For the first time, we characterized variation in mitochondrial density in relation to prenatal THs and temperature in birds, and in a wild population. In contrast to our predictions, we did not observe clear context-dependent effects of THs on mtDNA copy number (hormone × heating, F_1,155.8_=0.55, p = 0.458, Fig 1d, Table S5), and also no apparent effect of prenatal TH elevation (F_1,31.0_=0.739, p=0.397, Table S5) or heating treatment (F_1,41.5_=0.003, p=0.958, Table S5) *per se*. While THs are known to affect mitochondrial biogenesis and function postnatally (25, 26), only one previous study to our knowledge has investigated the effects of prenatal THs on mitochondrial parameters in a hypothyroid rat model. This study found no clear effects of prenatal THs on mitochondrial traits (28). Furthermore, mitochondria are generally responsive to varying (both high and low) ambient temperature (reviewed in 27). The lack of effects of our heating experiment may potentially be explained by the timing of the measurements: mitochondrial density was measured 5 days after the heating treatment had ceased, and given that mitochondrial traits are very plastic (as already shown in this species, 23), the effect of the temperature treatment may have vanished by the time of measurement. Finally, both TH and temperature effects on mitochondrial density could be tissue-specific, and not visible in blood cells.

We predicted that any effects of the treatments on growth, circulating TH levels or mitochondria could lead to oxidative stress, due to the altered production of free radicals, and/or antioxidant defenses. In line with the results above, we found no evidence for context-dependent effects of prenatal THs and postnatal temperature on the endogenous antioxidant glutathione or oxidative damage to lipids at the end of the heating period (d8) or at 13 days of age (all F < 1.61, p > 0.21, Fig. 1e,f, Table S6, S7). These results support our previous findings where elevated prenatal THs did not appear to influence postnatal oxidative stress biomarkers in birds (16, 17). Yet, effects of prenatal THs on oxidative stress could be tissue-dependent and/or only visible during embryo development, which needs to be further tested.

In conclusion, we found no clear evidence for context-dependent effects of prenatal THs depending on the ambient early postnatal temperature, nor support for the hypothesis that higher TH transfer to eggs in cold conditions benefits offspring in cooler rearing conditions. Our study nevertheless suggests multiple avenues for further research on the potential context-dependence of maternal effects on offspring phenotype and the potential underlying physiological mechanisms. As maternal effect has been suggested to enable rapid adaptation to climate change by providing an additional source of phenotypic plasticity (5–7), it is important to understand their context-dependent effects.

## Supporting information

ESM

## Acknowledgements

We thank all field assistants, especially Lucas Bousseau and Thomas Rosille and Päivi Kotitalo for their great effort, Janina Stauffer for help with laboratory analysis and Tuija Koivisto for analyzing the brooding videos. Turku Biocenter Finland supported the facilities for thyroid hormone measurements.

## Author contribution

BYH, AS & SR designed the study and conducted the fieldwork. All authors contributed to data analysis. BYH conducted the final statistical analysis. AS, NCS, MC & SR conducted laboratory work. BYH, AS & SR wrote the manuscript, with input from TS, MC and NCS.

## Data accessibility

All data will be archived and available in Dryad upon acceptance. Data is included as supplementary files (xx-xx).

