## Supplementary material for "Testing for context-dependent effects of maternal thyroid hormones on offspring survival and physiology: an experimental approach manipulating temperature in a wild bird species": ESM

### Mitochondrial density and molecular sexing analyses by qPCR

We extracted DNA from red blood cell pellet using a standard salt extraction alcohol precipitation method (Aljanabi and Martinez 1997). Extracted DNA was diluted in elution buffer BE for DNA preservation. DNA concentration and quality (260/280 and 260/230 ratios) were checked with a ND-1000-Spectrophotometer (NanoDrop Technologies, Wilmington, USA), and DNA integrity was verified in 48 samples chosen randomly using gel electrophoresis (50 ng of DNA, 0.8 % agarose gel at 100 mV for 60 min) and DNA staining with Midori Green. Each sample was then diluted to a concentration of 1.2 ng/µl for subsequent qPCR analysis.

Relative mitochondrial DNA copy number (mtDNAcn, an index of mitochondrial density) was quantified using a real-time quantitative PCR (qPCR) assays previously used and validated in this species (Stier et al. 2019). This technique estimates relative mtDNAcn as the ratio between one mitochondrial gene and one single copy nuclear gene. Here, we used RAG1 as a SCG (verified as single copy using a BLAST analysis on the collared flycatcher *Ficedula albicollis* genome) and cytochrome oxidase subunit 2 (COI2) as a mitochondrial gene (verified as non-duplicated in the nuclear genome using a BLAST analysis on the collared flycatcher *Ficedula albicollis* genome). Forward and reverse RAG1 primers were 5'-GCAGATGAACTGGAGGCTATAA-3' and 5'-CAGCTGAGAAACGTGTTGATTC-3', respectively. Forward and reverse COI2 primers were 5'-GGAGACGACCAAGTCTACAATG-3'; 5'-TTTCCGAACCCTCCGATTATG-3', respectively. For the qPCR assays, the reactions were performed on a 384-QuantStudio™ 12K Flex Real-Time PCR System (Thermo Fisher), in a total volume of 12µL including 6ng of DNA, primers at a final concentration of 200nM and 6µL of Absolute Blue qPCR Mix SYBR Green low ROX (Thermo Scientific). RAG1 and COI2 reactions were performed in triplicates on the same plates (6 plates in total); the qPCR conditions were: 15 min at 95°C, followed by 35 cycles of 15 s at 95°C, 30 s at 60°C and 30s at 72°C. A DNA sample being a pool of DNA from 10 individuals was used as a reference sample and was included in triplicate on every plate. The efficiency of each amplicon was estimated from a standard curve of the reference sample ranging from 1.5 to 24ng. The mean reaction efficiencies were 101.7 ± 2.8% for RAG1, 95.5 ± 1.8% for COI2. The relative mtDNAcn of each sample was calculated as $(1+Ef_{COI2})^{\Delta Cq\ COI2}/(1+Ef_{RAG1})^{\Delta CqRAG1}$; *Ef* being the amplicon efficiency, and ΔCq the difference in Cq-values between the reference sample and the focal sample. Intra-plate technical repeatability based on triplicates was 0.93 (95% C.I. [0.91-0.94]; N = 262). Inter-plate technical repeatability based on a few samples repeated on different plates was 0.96 (95% C.I. [0.86-0.99]; N = 12). The use of mtDNAcn as an index of mitochondrial density has been criticized in human (Larsen et al. 2012), but we have previously shown good correlations between mtDNAcn and mitochondrial respiration rates in pied flycatcher adult females (Stier et al. 2019).

Chicks were molecularly sexed using a qPCR approach adapted from Ellegren et al. 1997 and Chang et al. 2008. Forward and reverse sexing primers were 5'- CACTACAGGGAAAACTGTAC-3' (2987F) and 5'-CCCCTTCAGGTTCTTTAAAA -3' (3112R), respectively. qPCR reactions were performed in a total volume of 12µL including 6ng of DNA, primers at a final concentration of 800nM and 6µL of SensiFAST™ SYBR® Lo-ROX Kit (Bioline). qPCR conditions were: 3 min at 95°C, followed by 40 cycles of 45 s at 95°C, 60 s at 52°C and 60s at 72°C, then followed by a melting curve analysis (95°C 60s, 45°C 50s, increase to 95°C at 0.1°C/s, 95°C 30s). Samples were run in duplicates in a single plate and 6 adults of known sex were included as positive controls. Sex was determined by looking at the dissociation curve, with two peaks indicating the presence of a Z and W chromosome (female), and one peak indicating the presence of only the Z chromosomes (male).

**Table S1**. Generalized linear mixed model (GLMM) on the effects of prenatal thyroid hormone elevation (TH/Control) and postnatal temperature elevation (heated/non-heated) on nestling fledging success. Nestlings that died before sufficient exposure to heating treatment (i.e. before d4, n=11) were excluded from the analysis. The model was fit by maximum likelihood approach using Laplace approximation with binomial distribution and logit link function. Brood size, date, initial mass (d2) were included as covariates, and nest of origin and rearing as random intercepts. Main effects have been reported from a model with no interactions.

| **Fledging success (n=236)** | | | | |
|---|---|---|---|---|
| **Random effects:** | | **Variance** | **Std. Dev.** | |
| **Nest of origin (n=56)** | Intercept | 0.731 | 0.855 | |
| **Nest of rearing (n=56)** | Intercept | 17.879 | 4.228 | |
| **Fixed factors** | **Estimate** | **SE** | **z** | **p** |
| **Intercept** | 3.624 | 3.705 | 0.978 | |
| **Hormone (TH)** | 0.392 | 0.765 | 0.513 | 0.608 |
| **Heating (Non-heated)** | 1.534 | 1.662 | 0.923 | 0.356 |
| **Brood size** | -1.624 | 0.764 | -2.127 | **0.033** |
| **Body mass** | 2.126 | 0.680 | 3.127 | **0.002** |
| **Date** | -3.491 | 1.271 | -2.747 | **0.006** |
| **Hormone × Heating** | 0.196 | 1.259 | 0.156 | 0.876 |

N = 76 CO-non-heated, 50 CO-heated, 58 TH-non-heated, 52 TH-heated.

**Table S2**. General linear mixed model (GLMM) on the effects of prenatal thyroid hormone elevation (TH/Control) and postnatal temperature elevation (heated/non-heated) on body mass at (a) day 2, prior to heating treatment; (b) d8, after heating treatment and (c) d13 (close to fledging) after hatching. Brood size, date, initial mass (d2) and cross-fostering status (cross-fostered or not) have been included as covariates, and nest of origin and rearing as random intercepts when applicable. Sex was determined at d13. Main effects have been reported from a model with no interactions. The significance tests were conducted using Kenward-Roger approximation on the degrees of freedom.

| | | | | | | |
|---|---|---|---|---|---|---|
| **(a) Day-2 body mass (n=248)** | | | | | | |
| **Random effects:** | | **Variance** | **Std. Dev.** | | | |
| **Nest of origin (n=58)** | Intercept | 0.326 | 0.571 | | | |
| **Residual** | | 0.339 | 0.583 | | | |
| **Fixed effects:** | **Estimate** | **Std. Error** | **t** | **df** | **F** | **p** |
| **Intercept** | -138.700 | 185.800 | -0.747 | | | |
| **Hormone (TH)** | -0.193 | 0.170 | -1.140 | 1, 54.23 | 1.299 | 0.259 |
| **Date** | 0.008 | 0.011 | 0.765 | 1, 55.74 | 0.584 | 0.448 |
| **(b) Day-8 body mass (n=209)** | | | | | | |
| **Random effects:** | | **Variance** | **Std. Dev.** | | | |
| **Nest of origin (n=54)** | Intercept | <0.001 | <0.001 | | | |
| **Nest of rearing (n=52)** | Intercept | 0.839 | 0.916 | | | |
| **Residual** | | 0.605 | 0.778 | | | |
| **Fixed factors** | **Estimate** | **SE** | **t** | **df** | **F** | **p** |
| **Intercept** | 1518.487 | 329.132 | 4.614 | | | |
| **Hormone (TH)** | -0.075 | 0.131 | -0.571 | 1, 21.90 | 0.317 | 0.579 |
| **Heating (Non-heated)** | -0.014 | 0.295 | -0.046 | 1, 46.80 | 0.002 | 0.963 |
| **Brood size** | -0.265 | 0.124 | -2.135 | 1, 50.36 | 4.543 | **0.038** |
| **Cross-foster (yes)** | 0.218 | 0.121 | 1.805 | 1, 152.50 | 3.224 | 0.075 |
| **Day-2 body mass** | 1.109 | 0.092 | 12.113 | 1, 112.55 | 137.660 | **<0.001** |
| **Date** | -0.085 | 0.019 | -4.587 | 1, 21.01 | 47.690 | **<0.001** |
| **Hormone × Heating** | 0.311 | 0.257 | 1.210 | 1, 172.68 | 1.432 | 0.233 |

Table S2. (cont.)

| **(c) Day-13 body mass (n=183)** | | | | | | |
|---|---|---|---|---|---|---|
| **Random effects:** | | **Variance** | **Std. Dev.** | | | |
| **Nest of origin (n=53)** | Intercept | 0.316 | 0.562 | | | |
| **Nest of rearing (n=48)** | Intercept | 0.766 | 0.875 | | | |
| **Residual** | | 0.610 | 0.781 | | | |
| **Fixed factors** | **Estimate** | **SE** | **t** | **df** | **F** | **p** |
| **Intercept** | 1813.456 | 370.056 | 4.900 | | | |
| **Hormone (TH)** | 0.173 | 0.227 | 0.763 | 1, 32.30 | 0.565 | 0.456 |
| **Heating (Non-heated)** | 0.041 | 0.307 | 0.133 | 1, 37.18 | 0.018 | 0.896 |
| **Brood size** | -0.502 | 0.133 | -3.785 | 1, 43.08 | 14.163 | **<0.001** |
| **Cross-foster (yes)** | -0.089 | 0.140 | -0.636 | 1, 133.01 | 0.399 | 0.529 |
| **Day-2 body mass** | 0.397 | 0.122 | 3.265 | 1, 149.50 | 10.231 | **0.002** |
| **Date** | -0.102 | 0.021 | -4.863 | 1, 50.97 | 23.582 | **<0.001** |
| **Sex (male)** | 0.274 | 0.142 | 1.933 | 1, 142.73 | 3.652 | 0.058 |
| **Hormone × Heating** | 0.442 | 0.304 | 1.457 | 1, 150.27 | 2.068 | 0.153 |
| **Hormone × Sex** | -0.104 | 0.281 | -0.370 | 1, 137.70 | 0.134 | 0.715 |
| **Heating × Sex** | 0.357 | 0.287 | 1.246 | 1, 144.58 | 1.515 | 0.220 |

Sample sizes: Day 2: n = 132 CO, 116 TH;
Day 8: n = 60 CO-non-heated, 47 CO-heated, 53 TH-non-heated, 49 TH-heated
Day 13: n = 55 CO-non-heated, 42 CO-heated, 45 TH-non-heated, 41 TH-heated

**Table S3**. General linear mixed model (GLMM) on the effects of prenatal thyroid hormone elevation (TH/Control) and postnatal temperature elevation (heated/non-heated) on tarsus length at (a) d8, after heating treatment and (b) at close to fledging (d13) after hatching. Brood size, date, initial size, cross-fostering status (cross-fostered or not) and measurer have been included as covariates, and nests of origin and rearing as random intercepts. Sex was determined at d13. Main effects have been reported from a model with no interactions. The significance tests were conducted using Kenward-Roger approximation on the degrees of freedom.

| | | | | | | |
|---|---|---|---|---|---|---|
| **(a) Day-8 tarsus length (n=209)** | | | | | | |
| **Random effects:** | | **Variance** | **Std. Dev.** | | | |
| **Nest of origin (n=54)** | Intercept | <0.001 | <0.001 | | | |
| **Nest of rearing (n=52)** | Intercept | 0.503 | 0.709 | | | |
| **Residual** | | 0.382 | 0.618 | | | |
| **Fixed factors** | **Estimate** | **SE** | **t** | **df** | **F** | **p** |
| **Intercept** | 895.106 | 257.290 | 3.479 | | | |
| **Hormone (TH)** | -0.067 | 0.104 | -0.642 | 1, 21.95 | 0.400 | 0.534 |
| **Heating (Non-heated)** | 0.001 | 0.230 | 0.006 | 1, 44.83 | <0.001 | 0.996 |
| **Brood size** | -0.084 | 0.097 | -0.861 | 1, 48.33 | 0.739 | 0.394 |
| **Cross-foster (yes)** | 0.105 | 0.096 | 1.092 | 1, 152.38 | 1.180 | 0.279 |
| **Day-2 body mass** | 0.855 | 0.073 | 11.746 | 1, 111.52 | 129.380 | **<0.001** |
| **Date** | -0.050 | 0.015 | -3.419 | 1, 45.77 | 11.662 | **0.001** |
| **Measurer (LB)** | 0.052 | 0.371 | 0.141 | 1, 70.94 | 1.238 | 0.296 |
| **Measurer (TR)** | 0.376 | 0.389 | 0.966 | | | |
| **Hormone × Heating** | 0.260 | 0.203 | 1.277 | 1, 170.63 | 1.596 | 0.208 |
| **(b) Day-13 tarsus length (n=183)** | | | | | | |
| **Random effects:** | | **Variance** | **Std. Dev.** | | | |
| **Nest of origin (n=53)** | Intercept | 0.053 | 0.231 | | | |
| **Nest of rearing (n=48)** | Intercept | 0.119 | 0.345 | | | |
| **Residual** | | 0.152 | 0.390 | | | |
| **Fixed factors** | **Estimate** | **SE** | **t** | **df** | **F** | **p** |
| **Intercept** | 587.604 | 154.375 | 3.806 | | | |
| **Hormone (TH)** | -0.043 | 0.101 | -0.428 | 1, 32.66 | 0.177 | 0.677 |
| **Heating (Non-heated)** | 0.003 | 0.131 | 0.024 | 1, 35.90 | 0.001 | 0.981 |
| **Brood size** | -0.073 | 0.057 | -1.290 | 1, 42.41 | 1.639 | 0.207 |
| **Cross-foster (yes)** | -0.069 | 0.069 | -1.008 | 1, 138.44 | 0.998 | 0.320 |
| **Day-2 body mass** | 0.219 | 0.058 | 3.739 | 1, 148.17 | 13.324 | **<0.001** |
| **Date** | -0.032 | 0.009 | -3. 687 | 1, 47.46 | 13.532 | **<0.001** |
| **Sex (male)** | -0.019 | 0.070 | -0.279 | 1, 146.80 | 0.076 | 0.784 |
| **Measurer (LB)** | 0.759 | 0.445 | 1.705 | 2, 51.67 | 1.489 | 0.235 |
| **Measurer (TR)** | 0.784 | 0.451 | 1.741 | | | |
| **Hormone × Heating** | 0.012 | 0.150 | 0.078 | 1, 157.73 | 0.006 | 0.939 |
| **Hormone × Sex** | -0.044 | 0.138 | -0.320 | 1, 143.34 | 0.100 | 0.753 |
| **Heating × Sex** | 0.127 | 0.140 | 0.909 | 1, 150.95 | 0.803 | 0.372 |

Sample sizes: Day 8: n = 60 CO-non-heated, 47 CO-heated, 53 TH-non-heated, 49 TH-heated
Day 13: n = 55 CO-non-heated, 42 CO-heated, 45 TH-non-heated, 41 TH-heated

**Table S4**. General linear mixed model (GLMM) on the effects of prenatal thyroid hormone elevation (TH/Control) and postnatal temperature elevation (heated/non-heated) on (a) plasma triiodothyronine (T3) and (b) thyroxine (T4) concentrations (pg/ml) in day-13 nestlings. Body mass, cross-fostering status (cross-fostered or not), nestling sex, and hormone extraction batch have been included as covariates, and nests of origin and rearing as random intercepts. Main effects have been reported from a model with no interactions. The significance tests were conducted using Kenward-Roger approximation on the degrees of freedom.

| | | | | | | |
|---|---|---|---|---|---|---|
| **(a) Plasma T3 concentration (n=74)** | | | | | | |
| **Random effects:** | | **Variance** | **Std. Dev.** | | | |
| **Nest of origin (n=43)** | Intercept | 0.031 | 0.176 | | | |
| **Nest of rearing (n=37)** | Intercept | <0.001 | <0.001 | | | |
| **Residual** | | 0.032 | 0.179 | | | |
| **Fixed factors** | **Estimate** | **SE** | **t** | **df** | **F** | **p** |
| **Intercept** | -0.924 | 0.493 | -1.877 | | | |
| **Hormone (TH)** | 0.171 | 0.134 | 1.275 | 1, 28.28 | 1.553 | 0.223 |
| **Heating (Non-heated)** | 0.183 | 0.126 | 1.443 | 1, 27.52 | 1.992 | 0.169 |
| **Body mass** | 0.196 | 0.035 | 5.554 | 1, 36.48 | 26.892 | **<0.001** |
| **Sex (male)** | 0.074 | 0.128 | 0.579 | 1, 63.18 | 0.299 | 0.587 |
| **Cross-foster (yes)** | -0.134 | 0.122 | -1.091 | 1, 32.44 | 1.146 | 0.292 |
| **Batch (B)** | -0.048 | 0.169 | -0.287 | 2, 24.03 | 0.079 | 0.924 |
| **Batch (C)** | -0.062 | 0.155 | -0.397 | | | |
| **Hormone × Heating** | -0.335 | 0.237 | -1.415 | 1, 37.28 | 1.896 | 0.177 |
| **Hormone × Sex** | -0.318 | 0.253 | -1.259 | 1, 64.04 | 1.443 | 0.234 |
| **(b) Plasma T4 concentration (n=74)** | | | | | | |
| **Random effects:** | | **Variance** | **Std. Dev.** | | | |
| **Nest of origin (n=43)** | Intercept | 5.894 | 2.428 | | | |
| **Nest of rearing (n=37)** | Intercept | <0.001 | 0.002 | | | |
| **Residual** | | 6.772 | 2.602 | | | |
| **Fixed factors** | **Estimate** | **SE** | **t** | **df** | **F** | **p** |
| **Intercept** | 3.569 | 3.083 | 1.157 | | | |
| **Hormone (TH)** | 1.027 | 1.001 | 1.026 | 1, 34.47 | 1.032 | 0.317 |
| **Heating (Non-heated)** | 0.701 | 0.722 | 0.970 | 1, 22.37 | 0.895 | 0.354 |
| **Body mass** | 0.563 | 0.219 | 2.573 | 1, 33.16 | 5.648 | **0.023** |
| **Sex (male)** | 0.203 | 0.777 | 0.261 | 1, 53.19 | 0.060 | 0.808 |
| **Cross-foster (yes)** | -0.669 | 0.684 | -0.979 | 1, 26.25 | 0.917 | 0.347 |
| **Batch (B)** | 1.438 | 1.220 | 1.178 | 2, 33.54 | 0.731 | 0.489 |
| **Batch (C)** | 0.278 | 1.177 | 0.236 | | | |
| **Hormone × Heating** | -0.242 | 1.389 | -0.174 | 1, 32.08 | 0.028 | 0.867 |
| **Hormone × Sex** | -1.222 | 1.583 | -0.772 | 1, 60.57 | 0.537 | 0.467 |
| **Heating × Sex** | -0.291 | 1.531 | -0.190 | 1, 57.16 | 0.032 | 0.858 |

Sample sizes: n = 21 CO-non-heated, 20 CO-heated, 17 TH-non-heated, 16 TH-heated

**Table S5**. General linear mixed model (GLMM) on the effects of prenatal thyroid hormone elevation (TH/Control) and postnatal temperature elevation (heated/non-heated) on mitochondria density, ie. mitochondria DNA copy number, in day-13 nestlings. Mitochondria DNA copy numbers were ln-transformed to ensure a normal residual distribution. Body mass, nestling sex and cross-fostering status (cross-fostered or not) have been included as covariates, and nests of origin and rearing as random intercepts. Main effects have been reported from a model with no interactions. The significance tests were conducted using Kenward-Roger approximation on the degrees of freedom.

| **Mitochondria DNA copy number (n=183)** | | | | | | |
|---|---|---|---|---|---|---|
| **Random effects:** | | **Variance** | **Std. Dev.** | | | |
| **Nest of origin (n=53)** | Intercept | 0.013 | 0.114 | | | |
| **Nest of rearing (n=48)** | Intercept | 0.077 | 0.277 | | | |
| **Residual** | | 0.055 | 0.234 | | | |
| **Fixed factors** | **Estimate** | **SE** | **t** | **df** | **F** | **p** |
| **Intercept** | 1.563 | 0.288 | 5.432 | | | |
| **Hormone (TH)** | 0.049 | 0.056 | 0.879 | 1, 31.01 | 0.739 | 0.397 |
| **Heating (Non-heated)** | 0.005 | 0.090 | 0.053 | 1, 41.52 | 0.003 | 0.958 |
| **Body mass** | 0.005 | 0.020 | 0.243 | 1, 167.41 | 0.056 | 0.813 |
| **Sex (male)** | 0.023 | 0.042 | 0.539 | 1, 149.95 | 0.283 | 0.596 |
| **Cross-foster (yes)** | -0.035 | 0.041 | -0.858 | 1, 135.85 | 0.724 | 0.396 |
| **Hormone × Heating** | -0.069 | 0.091 | -0.755 | 1, 155.76 | 0.554 | 0.458 |
| **Hormone × Sex** | 0.064 | 0.081 | 0.781 | 1, 138.81 | 0.598 | 0.441 |
| **Heating × Sex** | 0.059 | 0.084 | 0.702 | 1, 147.56 | 0.479 | 0.490 |

Sample sizes: n = 55 CO-non-heated, 44 CO-heated, 45 TH-non-heated, 39 TH-heated

**Table S6**. General linear mixed model (GLMM) on the effects of prenatal thyroid hormone elevation (TH/Control) and postnatal temperature elevation (heated/non-heated) on blood total gluthathione concentration (tGSH) at (a) d8 and (b) d13 after hatching. Body mass and cross-fostering status (cross-fostered or not) have been included as covariates, and nests of origin and rearing as random intercepts. Sex was determined at d13. Main effects have been reported from a model with no interactions. The significance tests were conducted using Kenward-Roger approximation on the degrees of freedom.

| | | | | | | |
|---|---|---|---|---|---|---|
| **(a) Day-8 tGSH (n=173)** | | | | | | |
| **Random effects:** | | **Variance** | **Std. Dev.** | | | |
| **Nest of origin (n=45)** | Intercept | 0.004 | 0.067 | | | |
| **Nest of rearing (n=44)** | Intercept | 0.008 | 0.089 | | | |
| **tGSH assay (n=4)** | Intercept | 0.009 | 0.093 | | | |
| **Residual** | | 0.108 | 0.328 | | | |
| **Fixed factors** | **Estimate** | **SE** | **t** | **df** | **F** | **p** |
| **Intercept** | 0.159 | 0.244 | 0.650 | | | |
| **Hormone (TH)** | -0.073 | 0.056 | -1.293 | 1, 30.90 | 1.606 | 0.215 |
| **Heating (Non-heated)** | -0.038 | 0.058 | -0.651 | 1, 29.87 | 0.415 | 0.524 |
| **Body mass** | -0.021 | 0.019 | -1.106 | 1, 110.11 | 1.137 | 0.289 |
| **Cross-foster (yes)** | 0.045 | 0.051 | 0.875 | 1, 134.61 | 0.751 | 0.388 |
| **Hormone × Heating** | -0.089 | 0.106 | -0.838 | 1, 155.85 | 0.673 | 0.413 |
| **(b) Day-13 tGSH (n=149)** | | | | | | |
| **Random effects:** | | **Variance** | **Std. Dev.** | | | |
| **Nest of origin (n=45)** | Intercept | <0.001 | <0.001 | | | |
| **Nest of rearing (n=39)** | Intercept | 0.020 | 0.141 | | | |
| **tGSH assay (n=4)** | Intercept | 0.023 | 0.153 | | | |
| **Residual** | | 0.032 | 0.179 | | | |
| **Fixed factors** | **Estimate** | **SE** | **t** | **df** | **F** | **p** |
| **Intercept** | -0.377 | 0.207 | -1.819 | | | |
| **Hormone (TH)** | 0.014 | 0.033 | 0.411 | 1, 21.12 | 0.161 | 0.692 |
| **Heating (Non-heated)** | -0.007 | 0.055 | -0.133 | 1, 32.39 | 0.018 | 0.895 |
| **Body mass** | 0.019 | 0.013 | 1.405 | 1, 108.91 | 1.813 | 0.181 |
| **Sex (male)** | 0.027 | 0.034 | 0.806 | 1, 129.07 | 0.623 | 0.432 |
| **Cross-foster (yes)** | 0.064 | 0.031 | 2.052 | 1, 110.91 | 4.114 | **0.045** |
| **Hormone × Heating** | -0.104 | 0.066 | -1.565 | 1, 125.20 | 2.367 | 0.127 |
| **Hormone × Sex** | 0.056 | 0.067 | 0.837 | 1, 127.42 | 0.669 | 0.415 |
| **Heating × Sex** | -0.027 | 0.068 | -0.409 | 1, 126.67 | 0.158 | 0.692 |

Sample sizes: Day 8: n = 52 CO-non-heated, 39 CO-heated, 45 TH-non-heated, 37 TH-heated
Day 13: n = 47 CO-non-heated, 34 CO-heated, 37 TH-non-heated, 31 TH-heated

**Table S7**. General linear mixed model (GLMM) on the effects of prenatal thyroid hormone elevation (TH/Control) and postnatal temperature elevation (heated/non-heated) on blood lipid peroxidation (malonaldehyde, MDA concentration) at (a) d8 and (b) d13 after hatching. Body mass and cross-fostering status (cross-fostered or not) have been included as covariates, and nests of origin and rearing as random intercepts. Sex was determined at d13. Main effects have been reported from a model with no interactions. The significance tests were conducted using Kenward-Roger approximation on the degrees of freedom.

| **(a) Day-8 MDA (n=172)** | | | | | | |
|---|---|---|---|---|---|---|
| **Random effects:** | | **Variance** | **Std. Dev.** | | | |
| **Nest of origin (n=45)** | Intercept | 0.003 | 0.056 | | | |
| **Nest of rearing (n=44)** | Intercept | 0.017 | 0.130 | | | |
| **Residual** | | 0.043 | 0.208 | | | |
| **Fixed factors** | **Estimate** | **SE** | **t** | **df** | **F** | **p** |
| **Intercept** | -2.607 | 0.182 | -14.347 | | | |
| **Hormone (TH)** | -0.011 | 0.039 | -0.293 | 1, 26.37 | 0.083 | 0.775 |
| **Heating (Non-heated)** | -0.005 | 0.052 | -0.102 | 1, 35.09 | 0.010 | 0.920 |
| **Body mass** | -0.039 | 0.014 | -2.721 | 1, 149.63 | 6.992 | **0.009** |
| **Cross-foster (yes)** | -0.062 | 0.033 | -1.888 | 1, 123.13 | 3.523 | 0.063 |
| **Hormone × Heating** | -0.044 | 0.071 | -0.624 | 1, 144.79 | 0.380 | 0.539 |
| **(b) Day-13 MDA (n=149)** | | | | | | |
| **Random effects:** | | **Variance** | **Std. Dev.** | | | |
| **Nest of origin (n=45)** | Intercept | 0.008 | 0.087 | | | |
| **Nest of rearing (n=39)** | Intercept | 0.024 | 0.156 | | | |
| **Residual** | | 0.016 | 0.126 | | | |
| **Fixed factors** | **Estimate** | **SE** | **t** | **df** | **F** | **p** |
| **Intercept** | -3.094 | 0.171 | -18.095 | | | |
| **Hormone (TH)** | -0.028 | 0.038 | -0.746 | 1, 26.28 | 0.544 | 0.467 |
| **Heating (Non-heated)** | -0.022 | 0.056 | -0.388 | 1, 31.19 | 0.150 | 0.702 |
| **Body mass** | <0.001 | 0.012 | -0.008 | 1, 141. 74 | <0.001 | 0.994 |
| **Sex (male)** | 0.005 | 0.027 | 0.188 | 1, 118.00 | 0.034 | 0.853 |
| **Cross-foster (yes)** | 0.033 | 0.023 | 1.436 | 1, 99.72 | 2.031 | 0.157 |
| **Hormone × Heating** | -0.094 | 0.052 | -1.817 | 1, 115.39 | 3.209 | 0.076 |
| **Hormone × Sex** | 0.020 | 0.052 | 0.387 | 1, 112.53 | 0.146 | 0.703 |
| **Heating × Sex** | -0.054 | 0.054 | -1.005 | 1, 120.32 | 0.976 | 0.325 |

Sample sizes: Day 8: n = 52 CO-non-heated, 39 CO-heated, 45 TH-non-heated, 36 TH-heated
Day 13: n = 47 CO-non-heated, 34 CO-heated, 37 TH-non-heated, 31 TH-heated

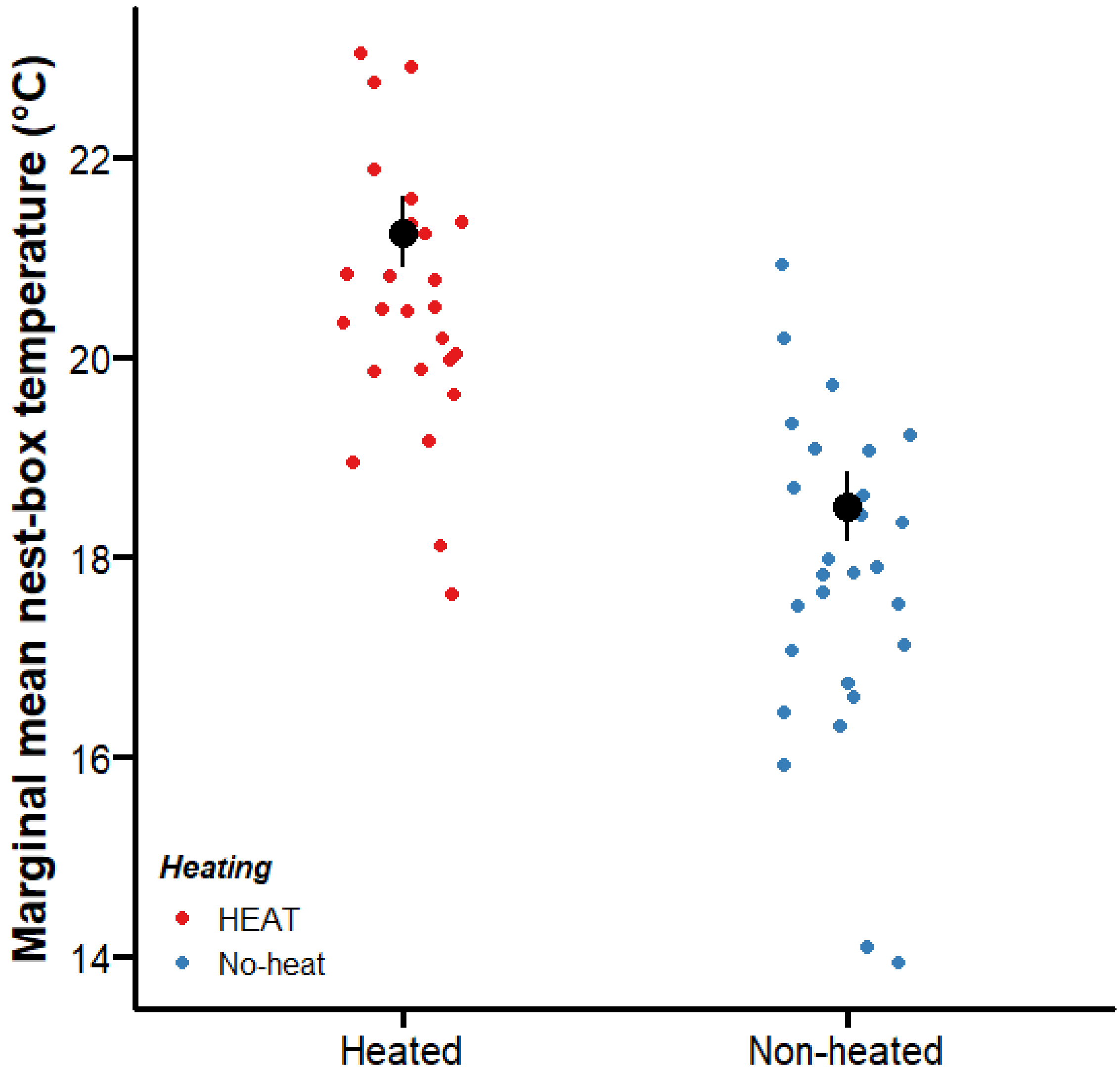


**Figure S1. Average nest-box temperatures across the period of heating treatment**. Small dots represent the average nest-box temperature from day 2 to day 8. Large black dots (mean±SE) represent marginal mean temperature controlled for date and iButton position.
